# Ongoing rapid evolution of a post-Y region revealed by chromosome-scale genome assembly of a hexaploid monoecious persimmon (*Diospyros kaki*)

**DOI:** 10.1101/2022.12.29.522208

**Authors:** Ayano Horiuchi, Kanae Masuda, Kenta Shirasawa, Noriyuki Onoue, Naoko Fujita, Koichiro Ushijima, Takashi Akagi

## Abstract

Sex chromosomes have evolved independently in many plant lineages. They have often undergone rapid structural degeneration and extension of non-recombining regions, which is conventionally considered to be strongly associated with the expression of sexually dimorphic traits. In this study, we assembled a monoecious hexaploid persimmon (*Diospyros kaki*) in which the Y chromosome had lost its function in male determination. Comparative genomic analysis among *D. kaki* and its dioecious relatives revealed that the non-functional Y chromosome (Y^*m*^) via silencing of the sex-determining gene, *OGI*, arose approximately two million years ago. Comparative analyses of the whole X and Y^*m*^ chromosomes suggested that the non-functional male-specific region of the Y-chromosome (MSY), or post-MSY, retained certain conserved characteristics of the original functional MSY. Specifically, comparison of the functional MSY in *D. lotus* and the non-functional post-MSY in *D. kaki* indicated that the post-MSY had been rapidly rearranged mainly via ongoing transposable element bursts, as well as in the functional MSY. These results suggest a novel interpretation that the rapid evolution of the post-MSY (and possibly also MSYs in dioecious *Diospyros* species) might reflect the ancestral genomic properties of these regions, rather than the evolution of male-determining functions and/or sexually dimorphic traits.

## Introduction

In contrast to animals, many different plant lineages have independently evolved chromosomal sex determination (or dioecious) systems from functionally hermaphrodite ancestors (Westergaard 1958, Charlesworth 1985, Ming et al. 2011, Henry et al. 2018). A comparative framework can therefore shed light on the diversity of routes by which sex determination and sex chromosomes may evolve. Recent advances in genomic technologies have revealed that the sex determining factors in different plant lineages differ in the molecular developmental functions involved, and also the evolutionary pathways that led to separate sexes (Akagi et al. 2014, 2018, 2019, Harkess et al. 2017, 2020, Muller et al. 2020, Kazama et al. 2022). Some species have single sex-determining genes or small sex-linked regions, while other plants have sex chromosomes, distinguished by the presence of physically extensive non-recombining regions (sometimes resulting from recombination suppression), and sometimes cytologically detectable heteromorphism. The loss of recombination may sometimes be associated with the evolution of sexual dimorphic phenotypic traits, which can lead to the establishment of sexually antagonistic polymorphisms (Charlesworth and Charlesworth 1978, Rice et al. 1992). Renner and Muller (2021) have questioned whether sex chromosome evolution in different plant lineages shares any common rules, and noted that the sizes of the male specific regions (Y-linked regions, or MSYs) are not related with the ages of different plant sex determining systems or the gene densities in the MSY, which might reflect the potential to be involved in the evolution of sexually dimorphic traits. In kiwifruit (the genus *Actinidia*), sexual dimorphisms being conserved across the genus. Nevertheless, transpositions of the two sex-determining factors have recurrently and independently formed new sex-linked regions with hemizygous MSYs (Akagi et al. unpublished). These MSYs contain only three conserved genes, including two sex-determining genes, one of which controls most of the sexual dimorphisms. This example suggests that MSYs might often evolve by a process different from the one involving the evolution of sexual dimorphism outlined above. One possibility proposes that a lack of recombination could be the ancestral state for some MSY regions, as plant (and animal) genomes often include extensive pericentromeric regions in which recombination is rare, and in some cases recombination is also rare in telomeric regions (Charlesworth 2019).

In persimmon (in the genus *Diospyros*), diploid species are dioecious, apart from hermaphroditic mutants/lines (Masuda and Akagi et al. 2022). The sexes are determined by a Y-linked gene, *OGI*, which formed by a duplication of an autosomal counterpart gene, *MeGI*, and expresses a small RNA that suppresses *MeGI* expression (Akagi et al. 2014). *MeGI* encodes a homeodomain ZIP1 (HD-ZIP1) which has neofunctionised via a lineage-specific duplication to gain roles in both suppressing androecium development and promoting gynoecium growth (Yang et al. 2019, Akagi et al. 2020). In contrast, individual trees of the hexaploid Oriental persimmon (*D. kaki;* 2n=6x=90) are mainly either gynoecious or wholly monoecious, with occasional production of hermaphroditic flowers on monoecious trees (Masuda et al. 2022, Masuda and Akagi 2022). Genetically female individuals (with hexaplex X: 6A+6X) are gynoecious, whereas genetically male individuals (carrying at least one Y chromosome) can be monoecious (Akagi et al. 2016a, Masuda et al. 2020). In the *D. kaki* Y-linked region, *OGI* was largely silenced by insertion of a SINE-like sequence named *Kali* into the promoter region (Akagi et al. 2016a). *MeGI* in this species has a novel epigenetic cis-regulatory developmental switch that controls its expression pattern, and can produce either male or female flowers (Akagi et al. 2016a, 2022). Thus *D. kaki* has lost the male-determining function, and become monoecious. Its Y acts as a Y^*m*^ factor, employing a similar terminology as Y^*h*^, which is used for the Y of hermaphrodite revertants of dioecious papaya (Wang et al. 2012).

Here, by chromosome-scale whole-genome assembly of monoecious *D. kaki* cv. Taishu (6A + XXXXYY; Akagi et al. 2016b), we clarify the history of Y^*m*^ and the evolution of the former MSY, by comparing Y^*m*^ with a functioning Y-linked region in a close diploid relative, the Caucasian persimmon (*D. lotus*).

## Results and Discussion

### Evolutionary paths of hexaploid persimmon and the silenced Y-determinant, *OGI^m^*

We assembled *Diospyros kaki* cv. Taishu (2n=6x=90, 6A + XXXXYY) whole genome sequences with PacBio HiFi reads, initially resulting in total 2.39Gbp scaffolds with N50 = 21.2Mbp (*N* = 37) (Supplementary Table S1-S2 for the basic genome characterization, Supplementary Fig. S1). Further scaffolding using RaGOO/RagTag (Alonge et al. 2019, 2022) with a reference genome of a close relative, *D. lotus* cv. Kunsenshi-male (2A + XY), and integration of allelic sequences resulted in generation of 14 chromosome scale autosomal scaffolds plus the XY pair (“pseudomolecule sequences”), consistent with the basic chromosome number of *Diospyros* species (*N* = 15). These scaffolds include 36,866 predicted genes covering 94.5% of the eudicot complete core gene set (complete BUSCOs (C)) (Figure 1B, Supplementary Table S1). All the genome sequences and the annotated data were deposited to the Persimmon Genome Database (http://persimmon.kazusa.or.jp/index.html) and Plant GARDEN (https://plantgarden.jp/en/index). Synteny analysis of the pseudomolecule sequences and analysis of the distribution of silent divergence (dS) values in putatively homologous gene pairs (see Materials and Methods), suggest that *D. kaki* underwent at least two paleo-genome duplication events, producing pairs with *dS* = 0.62-0.80 and 1.24-1.50, which would, respectively, be consistent with a *Diospyros-specific* genome duplication, Dd-α (Akagi et al. 2020) and the hexaploidization-γ that occurred in the common ancestor of eudicotyledonous plants (Jaillon et al. 2009) (Figure 1). Distributions of the *dS* values in *D. kaki* allelic sequences, and between orthologous gene pairs in *D. kaki* and its close diploid relatives, *D. lotus* (Akagi et al. 2020) or *D. oleifera* (Suo et al. 2020), (Figure 2A), suggest that the current diversity of hexasomic alleles in *D. kaki* (*dS* = 0.0141 for the peak) was established immediately after the divergence of *D. kaki* and *D. oleifera (dS* = 0.0225). The divergence of *D. lotus* may slightly predate these events, as *dS* is slightly larger (0.0301, *p* = 0.0013 for the *dS* distribution). Importantly, the pairwise *dS* values between the sequences of the SINE transposable element, *Kali*, within the *OGI* sequence (see above) in 12 cultivars from various East Asian area (Supplementary Table S3, Akagi et al. 2016b), range up to >0.02 (Figure 2A). These results suggest that this insertion coincided with the divergence of *D. kaki* and *D. oleifera*, and may predate the hexaploidization events, as summarized in Figure 2B. The *OGI* gene would then be estimated to have been established (creating a proto-Y) approximately 25 million years ago (mya), using an estimated rate of 4 × 10^−9^ substitutions per synonymous site per year in Arabidopsis (Beilstein et al. 2010, Wang et al. 2012), while the silence of *OGI (OGI^m^*) to become a Y^*m*^ factor in the *D. kaki* lineage occurred approximately 2 mya. Genome-wide synteny analysis with MCScanX supported the species divergence order estimated from the *dS* values, as *D. oleifera* exhibited more similar gene order to that in the *D. kaki* genome than to the *D. lotus* order (Figure 2C). Note, however, that this might be due to the incomplete pseudomolecules in the published *D. lotus* genome database (http://persimmon.kazusa.or.jp/). Genome-wide *dS* values between *D. kaki* and *D. oleifera* or *D. lotus* in 200-kb sliding windows were mostly consistent with the phylogenetic relationships of these species (Figure 2D). However, up to 3-4% of the genomic regions exhibited significantly lower *dS* values (*p* < 0.01 for each bin, Student’s *t*-test) between *D. kaki* and *D. lotus* than between *D. kaki* and *D. oleifera* (Figure 2E), suggesting potential introgressed regions from the *D. lotus* lineage, after the divergence of *D. kaki/D. oleifera* and *D. lotus*.

**Figure 1.**
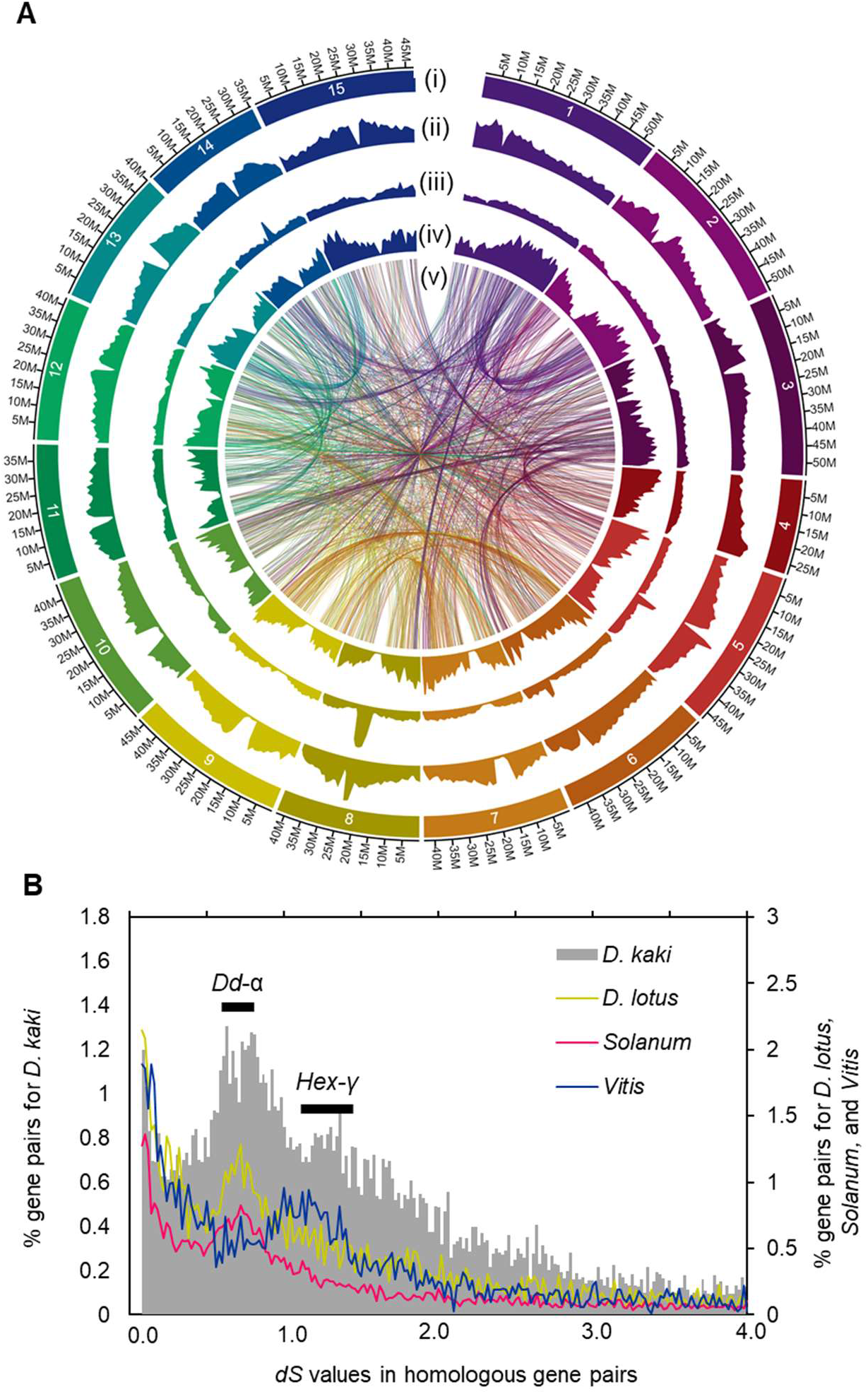
Characterization of *Diospyros kaki* cv. Taishu draft genome. **A.** Overview of 15 pseudomolecules comprising 14 autosomes (Chr1–14) and Y chromosome (Chr 15). From the outer layers inward, chromosome numbers (i), relative density of transposable elements detected by EDTA [(ii) for LTR-type, (iii) for non-LTR type], and relative gene density (iv), are given. In the central area (v), syntenic relationships within the persimmon genome, with gene pairs showing silent-site divergence (dS) = 0.6–0.9, which corresponds to a putative whole-genome duplication event, Dd-α (see panel **B**), are indicated. **B.** Distribution of *dS* rates between homologous gene pairs within the *D. kaki, D. lotus*, tomato (*Solanum lycopersicum*), and grape (*Vitis vinifera*) genomes. The *D. kaki* and *D. lotus* genomes show a consistent peak, corresponding to Dd-α, at the same *dS* value as the *Solanum* genome triplication. An additional peak in the *D. kaki* genome (dS = 1.24–1.50) is almost consistent with a peak in the *Vitis* genome, which would correspond to the hexaploidization y (Hex-Y) event.

**Figure 2.**
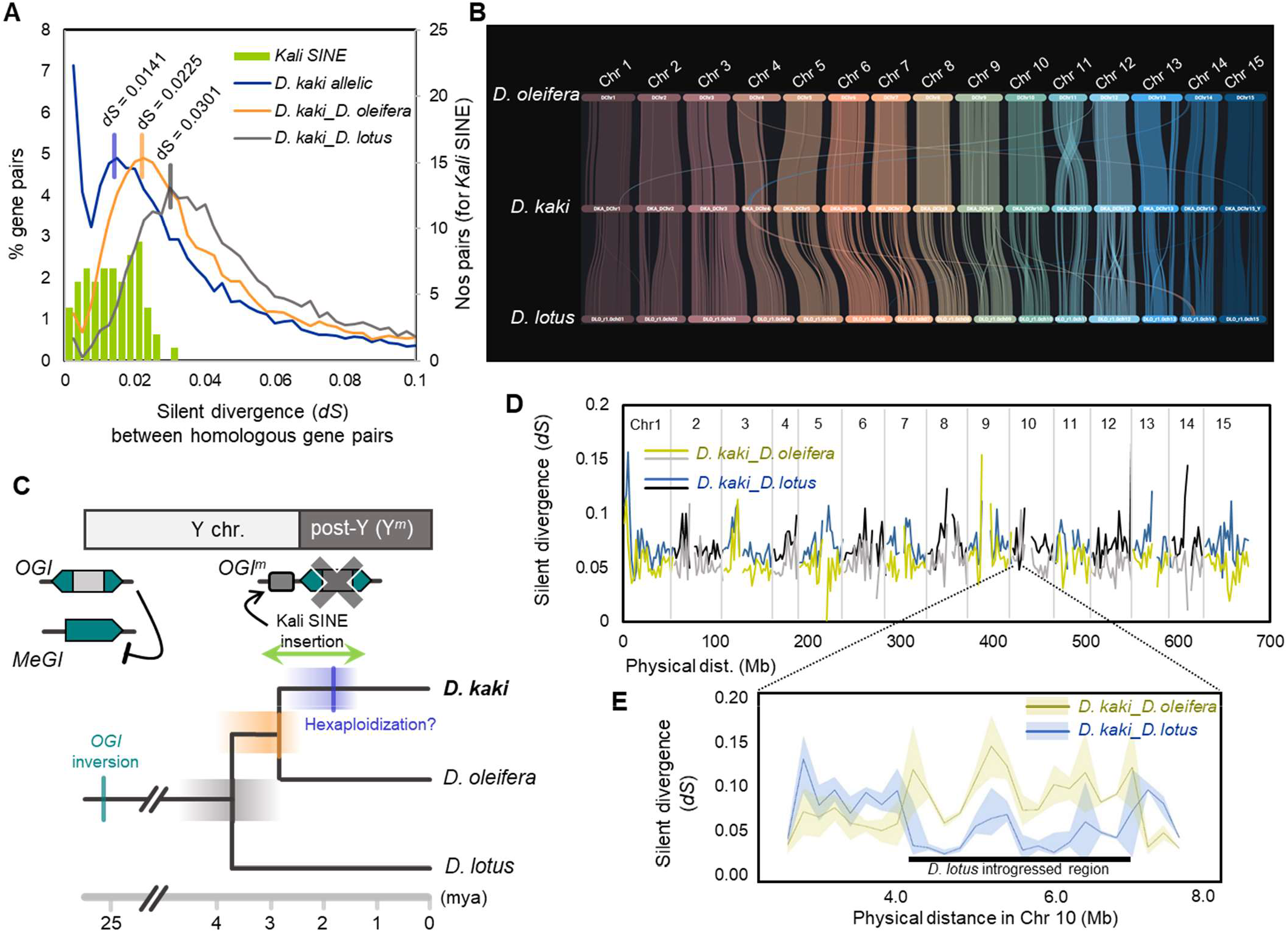
Species divergence and evolutionary path of the post-Y chromosome in *Diospyros kaki*. **A.** Comparison of the distribution of silent-site divergence (dS) values for *D. kaki* allelic sequences (blue), between the orthologs in *D. kaki* and *D. oleifera* (orange), and in *D. kaki* and *D. lotus* (gray). The distribution of *dS* values in the Kali-SINE allelic sequences is indicated by green bars. **B.** Chromosome-scale synteny analysis based on the gene orders in the *D. kaki, D. oleifera*, and *D. lotus* genomes. **C.** Schematic model for the evolution of the post-Y chromosome, including the *OGI^m^* silenced by the Kali-SINE insertion, with estimated ages. mya: million years ago. **D.** 1 Mb-bin walking analysis for the *dS* values between the orthologs in *D. kaki* and *D. oleifera*, and in *D. kaki* and *D. lotus*. Most of the genome showed smaller *dS* between *D. kaki* and *D. oleifera* than between *D. kaki* and *D. lotus*, which was consistent with the genome-wide *dS* distribution (shown in panel **A**). **E.** Expanded view of a region exhibiting smaller *dS* between *D. kaki* and *D. lotus* than between *D. kaki* and *D. oleifera*, implying potential introgression from the *D. lotus* genome.

### Conservation of MSY characteristics in Y^*m*^

Synteny analysis indicated highly conserved gene orders between the *D. kaki* X and Y^*m*^ chromosomes, except in the putative MSY (or post-MSY) region that includes the silenced *OGI^m^* (Figure 3A). The post-MSY occupies approximately 1.5Mbp in the pericentromeric region of chromosome 15 (Figure 3A, B, Supplementary Figure S2), with fragmented syntenic blocks compared with its X counterpart (Figure 3B). The *dS* values between X and Y^*m*^ alleles in monoecious *D. kaki* (Figure 3C) reached 0.212 in the central region flanking the *OGI* gene (Figure 3D), which is comparable to the *dS* values in the functional MSY of dioecious *D. lotus (dS* = 0.196 for the locus closest to OGI; Akagi et al. 2020), and the *dS* value between *OGI* and *MeGI* (*dS* = 0.205) corresponding to the initial establishment of the functional MSY (Akagi et al. 2014). We did not detect clear evidence for the formation of evolutionary strata, mainly due to the few X-Y allelic genes in the post-MSY, although the *dS* values between X- and Y-linked sequences decline with increased distance from *OGI^m^* (Figure 3D). This situation was consistent with the functional MSY in dioecious *D. lotus* (Akagi et al. 2020). These results suggest that the post-MSY in Y^*m*^ chromosome has conserved some characteristics of the original functional MSY.

**Figure 3.**
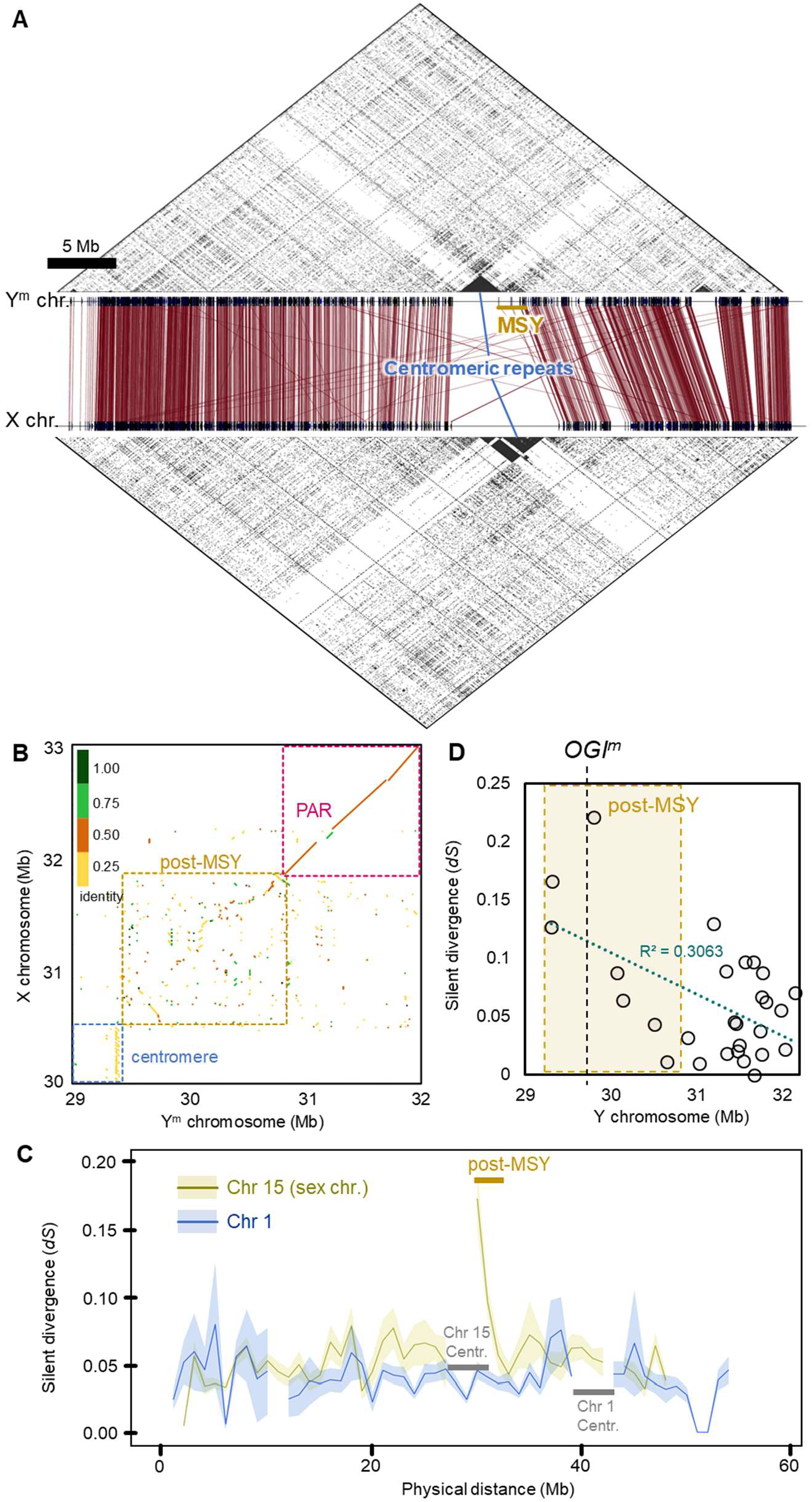
Comparative analysis of X and Y chromosomes in *Diospyros kaki*. **A.** Synteny based on the allelic genes between X and Y chromosomes. Putative allelic genes with significant synteny (<1e^−50^ in BLASTP) are connected with dark red lines. The upper and lower triangular areas indicate self-synteny dot plots. The post-MSY is located flanking the putative centromeric region. **B.** Synteny between the post-MSY and the counterpart X-allelic region. In the post-MSY, fragmental synteny blocks are observed. **C.** Transition of 1 Mb bin silent-site divergence (dS) values between X and Y allelic genes (dark yellow lines, with SD). As a control, the transition of 1 Mb bin *dS* values amongst the alleles for Chr 1 (or an autosome) is shown (blue lines, with SD). The post-MSY exhibited a distinct increase in *dS* values against the X alleles. **D.** The distribution of the *dS* values in each gene, in the 3 Mb region surrounding the post-MSY.

Under the two-factor model of the evolution of separate sexes and Y-linked regions (Charlesworth and Charlesworth 1978), hermaphrodite revertant individuals can arise, carrying so-called *“Y^h^”* chromosomes, and may have been favoured by artificial selection, as in the domestication of papaya (Wang et al. 2012, Van Buren et al. 2015, 2016) or in the grapevine (Zhou et al. 2017, 2019). To test the possibility that the persimmon Y^*m*^ has also experienced such selection, we analysed nucleotide diversity (*π*) and Tajima’s *D* values in the Y^*m*^ chromosome, using the genome-wide SNPs in resequencing data of 58 *D. kaki* cultivars (Supplementary Table S3, DRA015334 for the sequenced accessions in DDBJ). Recent selection for the Y^*m*^ allele should cause a “selective sweep”, with decreased diversity in the region. However, we did not detect this (Supplementary Fig. S3). This result would be consistent with the fact that all *D. kaki* Y chromosomes so far studied carry *OGI^m^* (Akagi et al. 2016a), and suggested that the establishment of Y^*m*^ could have been favoured by a strong naturally occurring bottleneck in population size longer ago, rather than by artificial selection, perhaps when the species became hexaploid, with Y^*m*^ maintained neutrally in domesticated *D. kaki*.

### Ongoing rapid evolution of the post-MSY

Despite only *ca*. 3.5 million years having elapsed since the divergence of *D. lotus* and *D. kaki* (see Figure 2B), their MSY regions differ by many rearrangements (Figure 4A, Supplementary Fig. S4A). Overall, only 5 of the 44 genes in the post-MSY are shared with the *D. lotus* MSY, and their physical orders were inverted (Supplementary Fig. S4B), in contrast to the flanking PAR, where most of the genes were shared between *D. lotus* and *D. kaki* (Supplementary Fig. S4C). The *D. kaki* post-MSY has also accumulated a high density of repetitive sequences, compared with the flanking PAR (Figure 4B). The *D. kaki* post-MSY, but not the corresponding region of the *D. kaki* X chromosome, is enriched with LTR-type transposable elements (TEs), especially the *Gypsy* class, unlike the *D. lotus* MSY (Figure 4C, Supplementary Fig. S5). The *D. lotus* MSY and the *D. kaki* post-MSY independently accumulated at least 31 mostly small (N ≧ 3) clusters of *Gypsy* class TEs (Supplementary Table S4). Approximately 2/3 of the *Gypsy* TEs in the post-MSY were not in clusters (using the criterion of >80% identity), suggesting that most are independently derived from sources elsewhere in the *D. kaki* genome. However, the largest *Gypsy* cluster (clst. 23) is a burst specific to the *D. kaki* post-MSY (Figure 4D). This cluster probably evolved very recently, after the establishment of the Y^*m*^, as pairwise *dS* values never exceed 0.02. Similar recent duplications of specific *Gypsy* TEs are also detected in the *D. lotus* MSY, but on a smaller scale (Supplementary Fig. S6).

**Figure 4.**
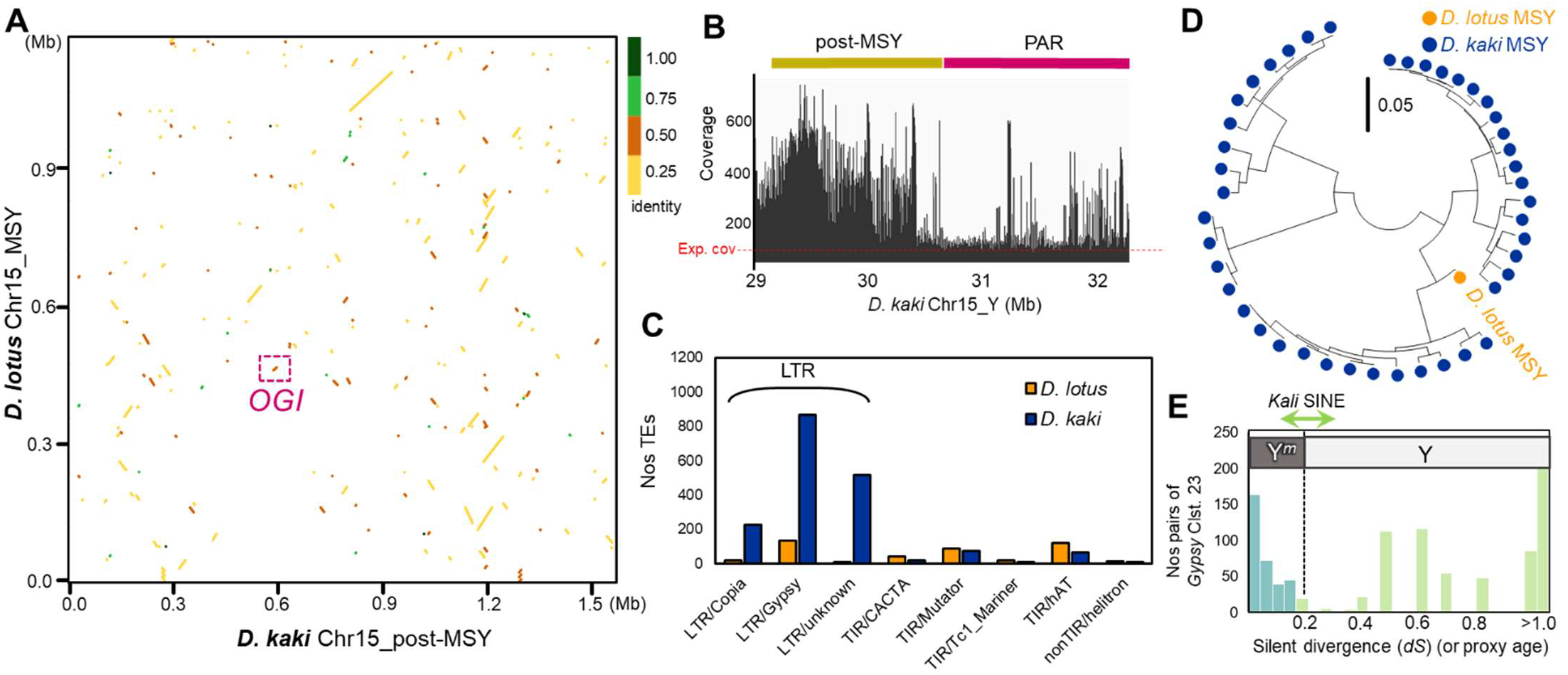
Comparative analysis of the MSY in *Diospyros lotus* and the post-MSY in *D. kaki*. **A.** Synteny between the MSY in *D. lotus* and the post-MSY in *D. kaki*. During the 3–4 million years since their divergence (see Fig. 1C), their structures have been highly rearranged. **B.** Mapping of the random gDNAseq data of *D. kaki* cv. Taishu to the post-MSY and the flanking PAR. The sequence coverage suggested that the post-MSY in *D. kaki* was enriched with repetitive sequences. **C.** Comparison of transposable elements (TEs) accumulation in the *D. lotus* MSY and the *D. kaki* post-MSY. The LTR-type TEs were more highly enriched in the *D. kaki* post-MSY. **D.** Phylogenetic analysis of a *Gypsy* cluster 23, which exhibited the *D. kaki* post-MSY-specific recent burst, with substitution rates < 0.05. **D.** Histogram of pairwise *dS* values in the *Gypsy* cluster 23, suggesting that dynamic TE bursts occurred after the Kali-SINE insertion, or establishment of the Y^*m*^ chromosome (or post-MSY).

**Figure 5.**
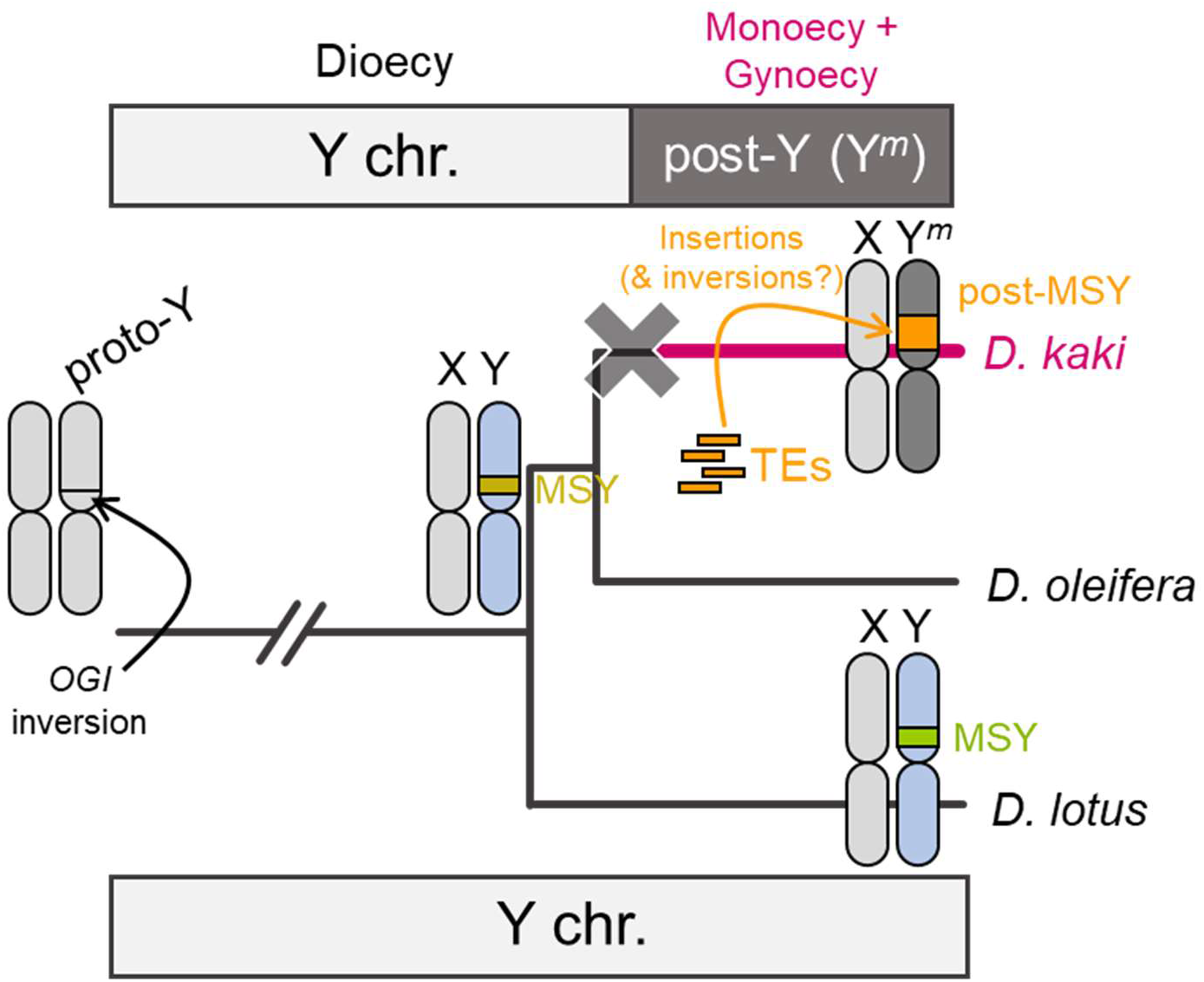
Model for the ongoing evolution of the post-Y chromosome. Since the establishment of the *OGI* inversion, the common ancestor of *D. lotus* and *D. kaki* had formed a putative MSY. Immediately after the divergence of *D. lotus* and *D. kaki, OGI* was silenced to establish the post-Y chromosome in the *D. kaki* ancestral lineage. The post-MSY has been continuously extended and rearranged via active TEs, which might be more rapid than for the MSY in dioecious *D. lotus*.

The highly rearranged structures in the *D. lotus* MSY and *D. kaki* post-MSY could reflect canonical MSY evolution, as rearrangements and TE insertions in non-recombining regions are not selectively disadvantageous (because crossing over does not occur, which also prevents ectopic exchanges between different TE insertions). MSYs in other plant species indeed show such changes, compared with their X counterparts (Akagi et al. 2019 for kiwifruit, Harkess et al. 2020 for garden asparagus, Ma et al. 2022 for spinach, Akagi et al. unpublished for the genus *Actinidia*). In *D. lotus*, selection may prevent some of these changes, because the MSY contains a functional OGI gene, and possibly other factors controlling sexual dimorphisms, but the post-MSY in *D. kaki*, might be under weak, or no, purifying selection, allowing insertions of TEs and duplications to occur. In monoecious *D. kaki* trees, morphological differences between males and females, such as inflorescence structure or different flower numbers (Supplementary Fig. S7), are similar to the sexual dimorphisms in diploid species in the genus *Diospyros*, and must thus be independent of the Y-linked region, or might be pleiotropic effect(s) of the sex determining gene(s). This possibility was originally suggested by Darwin (1877), who used the term “compensation”, and it is supported in persimmon (in the genus *Diospyros*) and kiwifruit (in the genus *Actinidia*) (Akagi and Charlesworth 2019). An interesting observation is that, in both these genera, their extended MSYs are located within ancestrally repetitive (and therefore probably rarely recombining) pericentromeric or peritelomeric regions (Akagi et al. unpublished). Hence, the recent evolution of the post-MSY (and possibly also MSYs in dioecious species) might reflect these regions’ ancestral genomic properties, rather than having evolved male-determining functions and/or sexually dimorphic traits.

## Materials and Methods

### Genome assembly

For whole-genome assembly, *D. kaki* cv. Taishu young leaves were sampled at the Grape and Persimmon Research Station, Institute of Fruit Tree and Tea Science, National Agriculture and Food Research Organization (NARO), Higashihiroshima, Japan. Genomic DNA for sequencing was extracted using the Genome-tip G100 (Qiagen, Venlo, Nederland). The genomic DNA was sheared into ~20 kb DNA fragments with a g-TUBE Prep Station (Covaris, Woburn, MA, USA) and a HiFi SMRTbell library was constructed with the SMRTbell Express Template Prep Kit 2.0 (PacBio, Menlo Park, CA, USA). The library DNA was fractionated using a BluePippin (Sage Science, Beverly, MA, USA) to eliminate fragments <20 kb and sequenced with a Four SMRT Cell 8M on the Sequel II system (PacBio). The sequence reads were converted into HiFi reads with the ccs pipeline (PacBio; https://ccs.how) and assembled with Hifiasm (Cheng et al. 2021). The obtained contigs were aligned to a reference genome sequence of the diploid male *D. lotus* ‘Kunsenshi-male’ (Akagi et al. 2020; 2n = 2x = 30, 2A+XY) for scaffolding using RaGOO/RagTag (Alonge et al. 2019, 2022) to build pseudomolecule sequences, with manual revisions as necessary.

### Repeat and gene annotations

Repetitive sequences in the assemblies were identified with phRAIDER (Schaeffer et al. 2016) and RepeatMasker (Smit et al. 2015; https://www.repeatmasker.org/) using *de novo* repeat libraries built with RepeatModeler2 (Flynn et al. 2020). Repeat elements were classified into nine types with RepeatMasker: short interspersed nuclear elements (SINEs), long interspersed nuclear elements (LINEs), long terminal repeat (LTR) elements, DNA elements, small RNA, satellites, simple repeats, low-complexity repeats, and unclassified.

Protein-coding genes were predicted from the repeat-masked genome sequences. Gene prediction was conducted with the Braker2 pipeline, trained with 22 Illumina short-reads mRNA-seq data from a variety of organs (fruit flesh, Maeda et al. 2019; flower buds and flowers, Masuda et al. 2022; and young leaf/flower primordia, Akagi et al. 2016a), at various developmental stages, in accordance with a previous pipeline (Shirasawa et al. 2022). The completeness of the assemblies was assessed using the BUSCO score (Simao et al. 2015).

### Synteny analyses

Chromosome-scale sequence synteny was evaluated with D-GENIES (Cabanettes and Klopp 2018) for dot-plot visualization (http://dgenies.toulouse.inra.fr/). Two whole-genome sequence (fasta) files were aligned with Minimap2 using the D-GENIES default settings. Large-scale synteny based on the gene orders was examined with MCScanX (Wang et al. 2012), in which the detected collinearity was visualized using SynVisio (https://synvisio.github.io/#/). All-versus-all BLASTP analyses were performed among the protein sequences in the *D. lotus, D. oleifera*, and *D. kaki* genomes, with an e-value cut-off of <1e^−40^. Syntenic blocks were constructed using MCScanX, with BLASTP and gff files, after preprocessing to be suitable for MCScanX. Intragenomic collinearity was evaluated by all-versus-all BLASTP analyses (<e^−50^ in BLASTP), for the whole genes, followed by selection with threshold values for silent-site divergence (dS; described later) and visualization with Circa software (https://omgenomics.com/circa/). Short-range syntenic blocks (or repetitive blocks within a chromosome) were identified with MUMmer4 (Marçais et al. 2018), using the nucmer program with the --maxmatch argument (with minimum length of the syntenic block = 25 bp).

### Detection of genetic diversity and age estimation

For detection of genetic diversity within the paralogs in the interspecific comparisons, gene pairs between *D. kaki* and *D. lotus*, or *D. kaki* and *D. oleifera*, that exhibited significant sequence similarity (<e^−50^ in BLASTP) were subjected to in-codon frame alignment using their protein and nucleotide sequences with Pal2Nal and MAFFT ver. 7 under the L-INS-i model (Katoh and Standley 2013). The *dS* values (with Jukes–Cantor correction) for the alignments were estimated with MEGA X (Kumar et al. 2018). For genetic diversity within the genome-wide *D. kaki* alleles, the whole genes in the pseudomolecule sequences (14+XY chromosomes) were aligned to the predicted genes in the initial scaffolds excluding the components of the pseudomolecule sequences, followed by detection of *dS* values, as described. For detection of *dS* values surrounding the *Kali* SINE, we sequenced the <2 kb PCR products flanking the *Kali* SINE (Akagi et al. 2016a, 2016b) and the mutated *OGI* in 12 cultivars (Supplementary Table S5). To estimate the divergence time between the gene pairs, we adopted an estimated rate of 4 × 10^−9^ substitutions per synonymous site per year in accordance with previous reports (Beilstein et al. 2010, Wang et al. 2012).

### Characterization of the genomic context around the MSY

For construction of Illumina genomic libraries, we used approximately 0.5 μg genomic DNA of *D. kaki* cv. Taishu. The libraries were constructed using the KAPA HyperPlus Library Preparation Kit (KAPA Biosystems) and sequenced using the Illumina HiSeq 4000 platform (with 150 bp paired-end reads). All Illumina sequencing was conducted at the Vincent J. Coates Genomics Sequencing Laboratory at the University of California, Berkeley. The raw reads were processed using custom Python scripts developed in the Comai laboratory and available online (http://comailab.genomecenter.ucdavis.edu/index.php/), as previously described (Akagi et al. 2014). The preprocessed reads were aligned to the whole-genomic scaffolds of each species, with the Burrows–Wheeler Aligner Maximal Exact Match algorithm (Li and Durbin 2009), allowing up to 12% mismatches. The mapped reads were visualized with Integrative Genomics Viewer (Robinson et al. 2011).

### *De novo* transposable element annotations and evolution

Transposable elements in the genomes of *D. kaki* and *D. lotus* were detected with the Extensive *denovo* TE Annotator (EDTA) pipeline, which integrates structure- and homology-based approaches for TE identification, including LTRharvest, LTR_FINDER_parallel, LTR_retriever, Generic Repeat Finder, TIR-Learner, MITE-Hunter, and HelitronScanner, with extra basic and advanced filters (Ou et al. 2019). Clustering of TEs within the *Gypsy* family was conducted with cd-hit-est in CD-HIT (Cluster Database at High Identity with Tolerance) (Li and Godzik 2006), with the -c 0.8 (>80% sequence identity) option.

To clarify the evolutionary topology of certain *Gypsy* clusters, we aligned their sequences with MAFFT ver. 7 with the L-INS-i model, followed by manual pruning using SeaView. The alignments were concatenated and all sites, including missing data and gaps, were used to construct phylogenetic trees with the maximum likelihood method using IQ-TREE 2 (Minh et al. 2020) with automatically optimized parameters.

### Accession numbers and construction of database

All genome sequences and the annotated data were deposited with the Persimmon Genome Database (http://persimmon.kazusa.or.jp/index.html) and Plant GARDEN (https://plantgarden.jp/en/index). All PacBio genome sequencing data were lodged with the DDBJ Sequence Read Archive (SRA) database (BioProject ID PRJDB14984).

## Acknowledgements

We thank Dr. Deborah Charlesworth (Institute of Evolutionary Biology, University of Edinburgh, UK) for discussion and comments for this study, and Edanz (https://jp.edanz.com/ac) for editing a partial draft of the manuscript. This work was supported by PRESTO from Japan Science and Technology Agency (JST) [JPMJPR20Q1 to T.A.] and Grant-in-Aid for Scientific Research (B) [22H02339 to T.A.] and for Transformative Research Areas (A) [22H05172 and 22H05173 to T.A., and 22H05181 to K.S.] from JSPS.

## Author contribution statement

T.A. conceived the study. A.H., K.M., and T.A. designed experiments. A.H., K.M., K.S., and T.A. conducted the experiments. A.H., K.M., K.S., and T.A. analyzed the data. N.O., N.F., K.U., and T.A. contributed to plant resources and facilities. A.H. and T.A. drafted the manuscript. All authors approved the manuscript.

