## Supplementary Information for "Ongoing rapid evolution of a post-Y region revealed by chromosome-scale genome assembly of a hexaploid monoecious persimmon (*Diospyros kaki*)"

**Supplementary Figure S1-S7**
**Supplementary Table S1-S4**

**Supplementary Fig. S1**

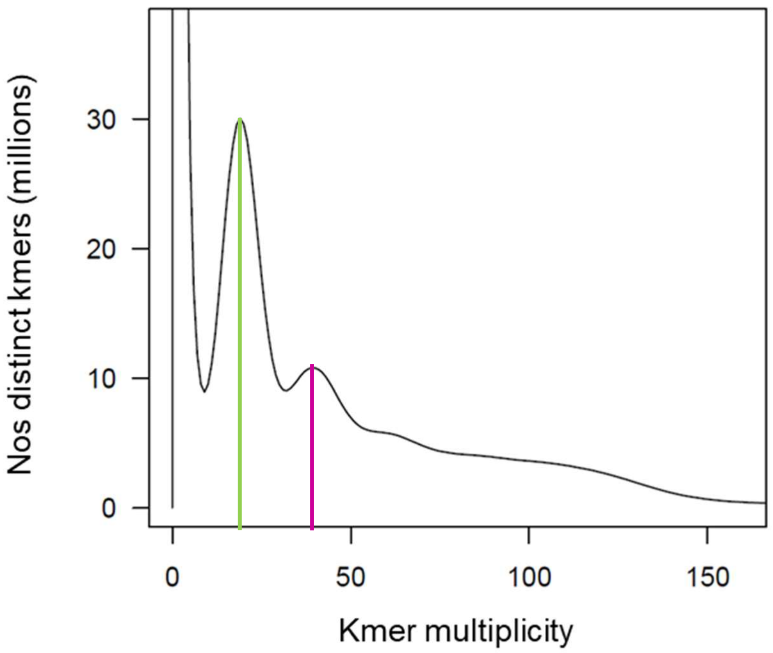


**Supplementary Fig. S1 kmer distribution analysis to estimate the degree of heterozygosity.**
The distribution of distinct k-mers (k = 17) from the Illumina short reads showed two peaks at multiplicities of 20 (light green) and 40 (magenta), which correspond represent heterozygous and homozygous sequences, respectively. Putative autohexaploidy alleles (or haploblocks) were frequently separately assembled, resulting in irregularly higher heterozygous sequences peak than homozygous sequences one.

**Supplementary Fig. S2**

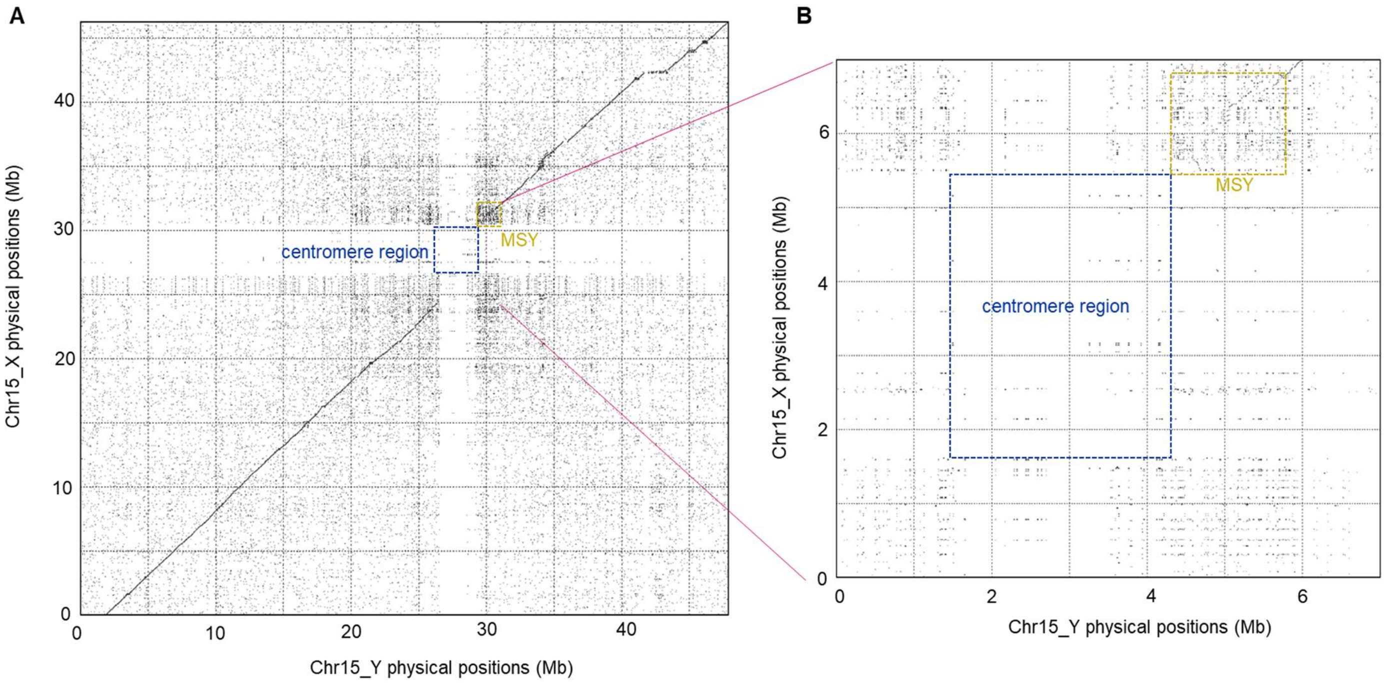


**Supplementary Fig. S2 Syntenic relationship in X and Y$^{m}$ chromosome**

Mummer plot synteny analysis between X and Y$^{m}$ chromosomes (**A**) and the closing up of the cenromeric/pericentromeric regions (**B**), suggested that the post-MSY locates on a pericentromeric region, ranging approximately 1.5Mbp. The putative centromere regions of X and Y chromosomes shared no substantial repeats (minimum length = 25bp per plot).

**Supplementary Fig. S3**

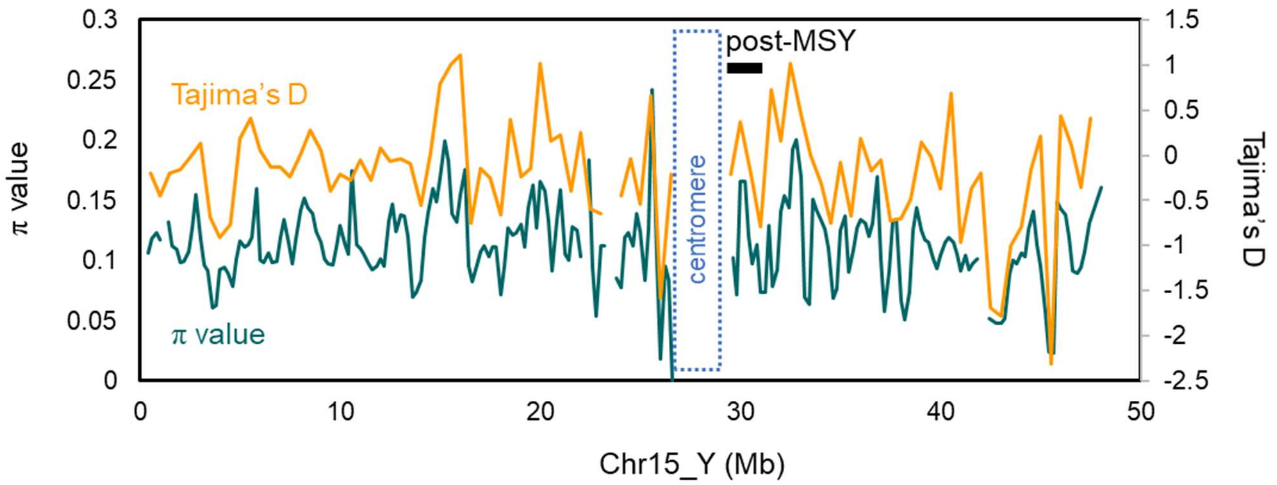


**Supplementary Fig. S3 Detection of potential selective sweep in Y*$^m$* chromosome**

We detected the transitions of nucleotide diversity (*π* values) and Tajima's *D* values in the Y$^m$ chromosome, using ddRAD-Seq data from 58 persimmon cultivars (DRA015334 for the sequence accessions in DDBJ). The Y$^m$ chromosome exhibited no substantial reduction in both *π* and Tajima's *D* values, except in *ca.* 45-Mbp region. This situation suggested no recent selection on the post-MSY.

**Supplementary Fig. S4**

A

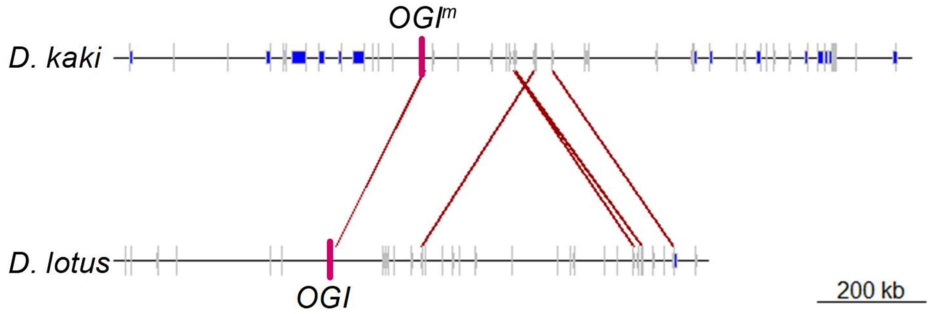


B

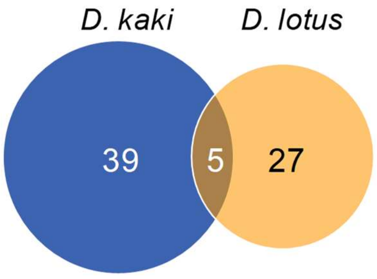


| *D. kaki* MSY gene ID | *D. lotus* MSY gene ID | putative gene functions |
|---|---|---|
| DKAch15y11764.t1 | Dlo_pri0560F.1_g00960.1 | Plant protein 1589 of unknown function |
| DKAch15y11756.t1 | Dlo_pri0792F.1_g00190.1 | unknown protein; LOCATED IN: plasma membrane |
| DKAch15y11753.t1 | Dlo_pri0792F.1_g00470.1 | Phosphatidylinositol N-acetyglucosaminlytransferase |
| DKAch15y11752.t1 | Dlo_pri0792F.1_g00480.1 | Disease resistance-responsive |
| DKAch15y11752.t1 | Dlo_pri0792F.1_g00510.1 | Disease resistance-responsive |

**Supplementary Fig. S4 Comparative analysis of the functional MSY in *D. lotus* and the non-functional post-MSY in *D. kaki***

Collinearity in gene orders between the MSY in *D. lotus* and the post-MSY in *D. kaki* (**A**). Putative orthologous genes were connected with red lines. Only 5 genes, including *OGI* (or $OGI^m$), showed significant similarity, while their physical orders/distances were highly rearranged. **B**, Venn diagram of the genes shared in the *D. kaki* post-MSY and the *D. lotus* MSY, and their functional annotations.

**Supplementary Fig. S5**

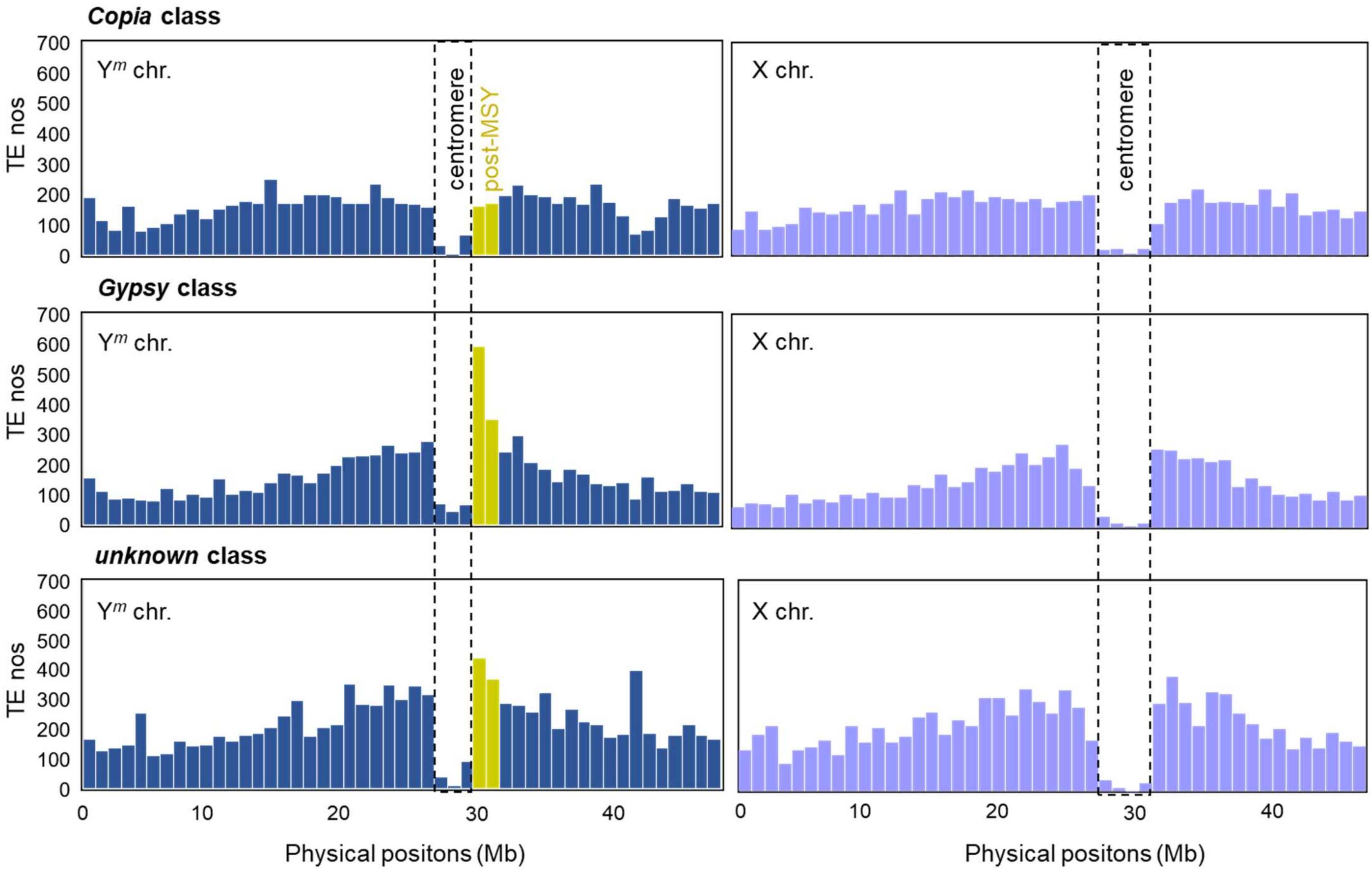


**Supplementary Fig. S5 Distribution of LTR-type TEs in X and Y$^m$ chromosomes in *D. kaki***

The *Copia*, *Gypsy* and *unknown* classes tended to be enriched in the post-MSY, in comparison to the counterpart X regions. Of them, the *Gypsy* class is particularly more accumulated in the pericentromeric post-MSY than in the counterpart X.

**Supplementary Fig. S6**

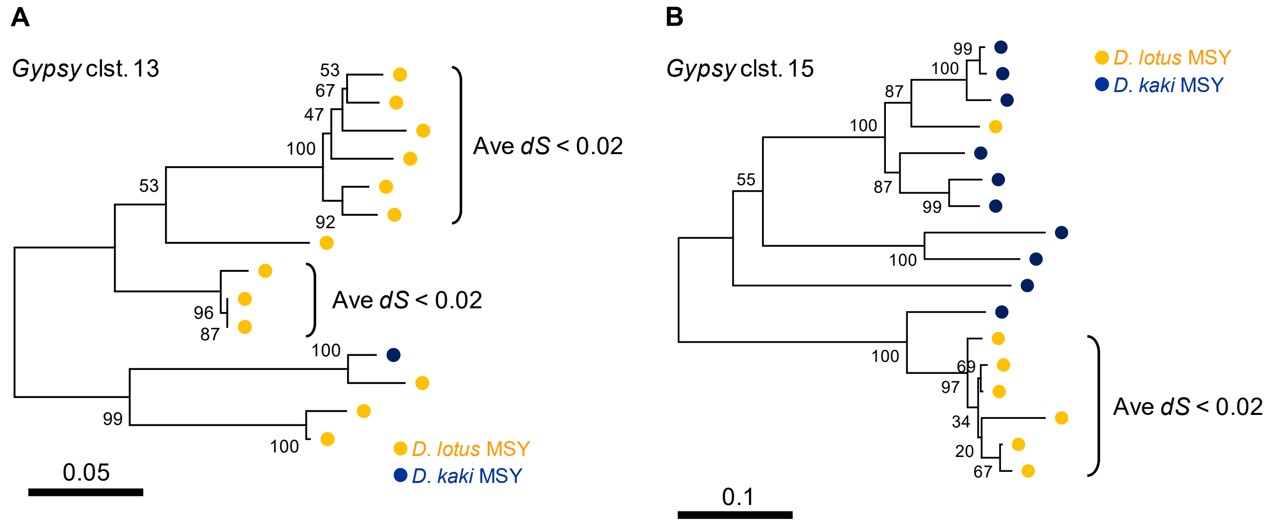


**Supplementary Fig. S6 Two *Gypsy* class clusters including recent duplications in the *D. lotus* MSY**

*Gypsy* clusters 13 (**A**) and 15 (**B**) (see Supplementary Table S4 for the details) showed *D. lotus*-specific duplications that putatively postdated the establishment of the $Y^m$ in *D. kaki* (or *dS* < 0.02). However, their scales were substantially smaller than the recent duplications specific to the post-MSY in *D. kaki*.

**Supplementary Fig. S7**

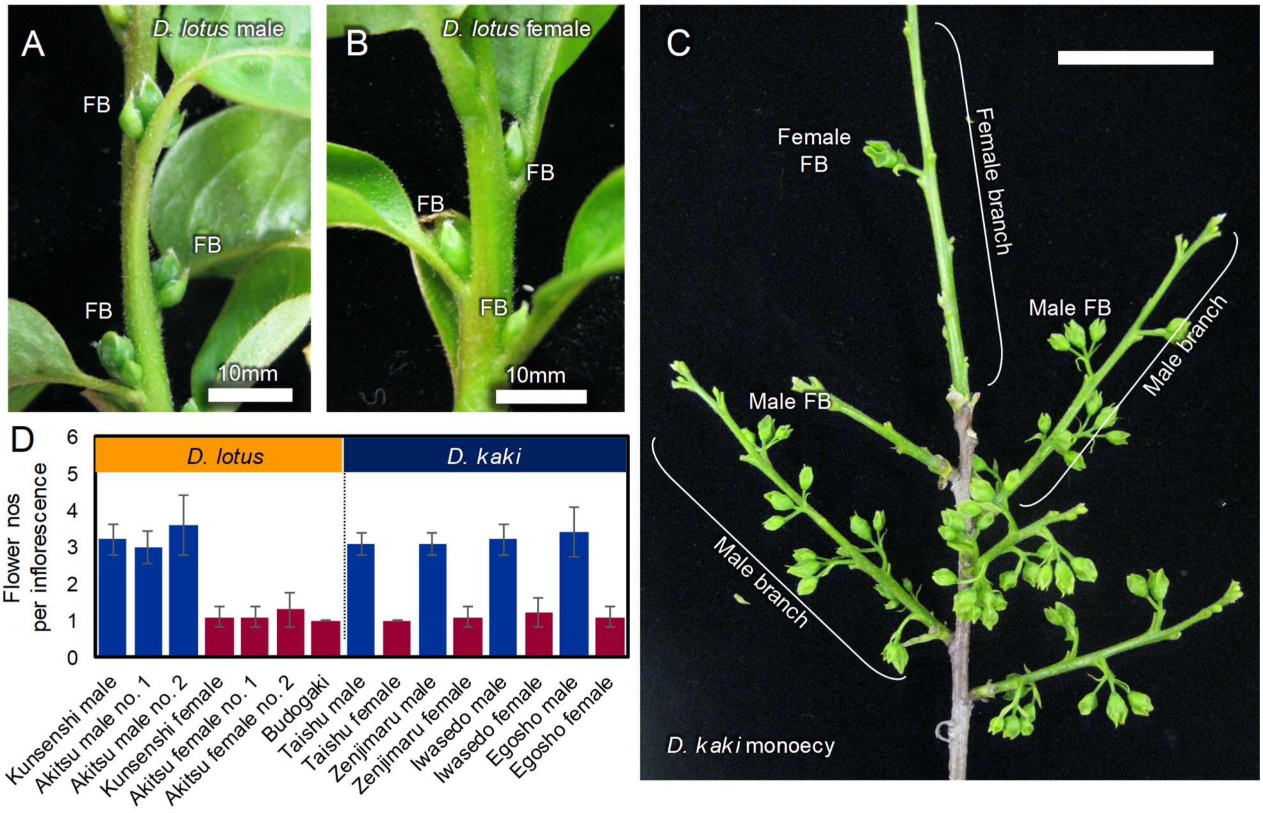


**Supplementary Fig. S7 Representative sexually dimorphic traits in *D. lotus* are shared with male/female flowers in monoecious *D. kaki***

**A-B**, *D. lotus* male individuals exhibit mostly trifurcated floral inflorescence (**A**), while female individuals exhibit mostly solitary inflorescence (**B**). FB: flower buds. This dimorphism is observed between male and female flowers in monoecious *D. kaki* (**C, D**). In *D. kaki*, each branch unit shows consistent sexuality (Akagi et al. 2016). Female branches carry statistically fewer flowers than male branches, which is also consistent with a sexually dimorphic trait in dioecious *D. lotus*.

### Supplementary Table S1 Basic characterization of the assembled *D. kaki* genome

| topic | initial assemble | pseudomolecule |
|---|---|---|
| **total length (bp)** | 2,394,259,740 | 708,766,281 |
| **contig nos** | 680 | 15 |
| **N50 (bp)** | 21,197,114 | 47,869,488 |
| **average length (bp)** | 3,520,970 | 47,251,085 |
| **largest contig (bp)** | 50,676,777 | 53,678,624 |
| **Complete BUSCOs (C)** | 99.3% | 94.5% |
| **Complete and single-copy (S)** | 6.3% | 86.2% |
| **Complete and duplicated (D)** | 93.0% | 8.3% |
| **Fragmented BUSCOs (F)** | 0.3% | 0.6% |
| **Missing BUSCOs (M)** | 0.4% | 4.9% |
| **Total BUSCO groups searched (nos)** | 1614 | 2326 |
| **Numbers of the genes** | | 36,866 |

### Supplementary Table S2 Distribution of transposable elements in the *D. kaki* genome.

| chromosome | Copia | Gypsy | unknown | CACTA | Mutator | Tc1_Mariner | hAT | PIF_Harbinger | Helitron |
|---|---|---|---|---|---|---|---|---|---|
| **Chr. 1** | 55,713 | 58,271 | 77,283 | 25,987 | 42,957 | 2,608 | 16,850 | 7,621 | 22,388 |
| **Chr. 2** | 7,513 | 8,089 | 10,624 | 3,455 | 5,984 | 344 | 2,381 | 1,104 | 3,030 |
| **Chr. 3** | 8,429 | 8,312 | 11,473 | 4,440 | 6,865 | 431 | 2,663 | 1,324 | 3,476 |
| **Chr. 4** | 4,491 | 4,427 | 6,339 | 2,135 | 3,140 | 165 | 1,409 | 649 | 1,732 |
| **Chr. 5** | 7,839 | 7,776 | 11,614 | 3,443 | 7,748 | 354 | 2,258 | 1,141 | 2,978 |
| **Chr. 6** | 6,713 | 7,782 | 10,595 | 3,259 | 6,802 | 361 | 2,342 | 941 | 2,902 |
| **Chr. 7** | 6,611 | 6,623 | 9,206 | 3,060 | 4,764 | 358 | 2,088 | 931 | 2,503 |
| **Chr. 8** | 6,424 | 9,033 | 12,251 | 3,221 | 13,957 | 322 | 2,052 | 898 | 2,720 |
| **Chr. 9** | 7,181 | 8,902 | 11,151 | 3,284 | 4,815 | 333 | 1,955 | 956 | 2,777 |
| **Chr. 10** | 6,960 | 6,774 | 9,477 | 3,339 | 5,152 | 354 | 2,138 | 937 | 2,815 |
| **Chr. 11** | 6,675 | 7,161 | 9,509 | 2,918 | 4,990 | 285 | 2,135 | 926 | 2,672 |
| **Chr. 12** | 7,027 | 7,122 | 8,815 | 3,023 | 5,229 | 316 | 2,017 | 940 | 2,856 |
| **Chr. 13** | 6,131 | 7,108 | 9,423 | 2,715 | 4,620 | 290 | 1,888 | 822 | 2,394 |
| **Chr. 14** | 5,970 | 6,636 | 8,889 | 3,039 | 5,076 | 286 | 1,601 | 824 | 2,157 |
| **Chr. 15-X** | 7,134 | 7,365 | 9,629 | 3,378 | 5,602 | 374 | 2,180 | 945 | 3,099 |
| **Chr. 15-Y** | 7,616 | 7,743 | 10,582 | 3,563 | 6,107 | 376 | 2,387 | 1,001 | 3,278 |

**Supplementary Table S3 List of plant materials**

| cultivar | putative origin | sample name in DDBJ | Detection of *dS* in the *OGI* promoter |
|---|---|---|---|
| Zenjimaru | Japan | libDk001 | X |
| Hazegosho | Japan | libDk004 | |
| Kakiyamagaki | Japan | libDk018 | |
| Egosho | Japan | libDk019 | X |
| Meotogaki | Japan | libDk021 | X |
| Taishu | Japan | libDk022 | X |
| Fujiwaragosho | Japan | libDk025 | X |
| Taiwanshoshi | Taiwan | libDk027 | X |
| Hanagosho | Japan | libDk030 | |
| Kanshu | Japan | libDk033 | |
| Nagara | Japan | libDk039 | |
| Kunitomi | Japan | libDk044 | X |
| Seihakuji | Japan | libDk047 | |
| Deshimaru | Japan | libDk049 | |
| Yamagaki | Japan | libDk054 | |
| Chagone | Japan | libDk066 | X |
| Kikumanjuu | Japan | libDk069 | |
| Yoshino | Japan | libDk072 | |
| Nishimurawase | Japan | libDk073 | X |
| Yashima | Japan | libDk076 | |
| Ibogaki | Japan | libDk083 | |
| Shogatsu | Japan | libDk087 | |
| Nitari | Japan | libDk088 | |
| Kawagone | Japan | libDk094 | |
| Akazu | Japan | libDk098 | |
| Saburoza | Japan | libDk101 | |
| Atagobou | Japan | libDk103 | |
| Shojo | Japan | libDk106 | |
| Sanenashi | Japan | libDk108 | |
| Kyara | Japan | libDk116 | |
| Tenjingosho | Japan | libDk126 | |
| Fudegaki | Japan | libDk129 | |
| Nanshi | Korea | libDk131 | X |
| Shirotodamashi | Japan | libDk141 | |
| Amayotsumizo | Japan | libDk145 | |
| Yotsumizo | Japan | libDk151 | |
| Yamatogosho | Japan | libDk152 | |
| Shoro | Japan | libDk160 | |
| Beniemon | Japan | libDk165 | |
| Mushirodagosho | Japan | libDk171 | |
| Shozaemon | Japan | libDk175 | |
| Iwasedo | Japan | libDk184 | |
| Ogosho | Japan | libDk187 | |
| Cal.Fuyu | Japan | libDk198 | |
| Suruga | Japan | libDk202 | |
| Monpei | Japan | libDk206 | |
| Muraya | Japan | libDk209 | |
| Beniwase | Japan | libDk210 | |
| Oniwa | Japan | libDk221 | |
| Emon | Japan | libDk222 | |
| Toyoka | Japan | libDk229 | |
| Shimokitahagakushi | Japan | libDk232 | |
| Tohachi | Japan | libDk238 | |
| Mikado | Japan | libDk240 | |
| Koudagosho | Japan | libDk246 | |
| Chichibuissaigaki | Japan | libDk254 | X |
| Saisho | Japan | libDk255 | |
| Okugosho | Japan | libDk259 | X |

**Supplementary Table S4 Clustering of *Gypsy* class TEs in the functional MSY of *D. lotus* and non-functional post-MSY in *D. kaki***

| Clust | *D. kaki* MSY | *D. lotus* MSY |
|---|---|---|
| **Cluster 0** | 3 | 0 |
| **Cluster 1** | 4 | 0 |
| **Cluster 2** | 7 | 1 |
| **Cluster 3** | 2 | 10 |
| **Cluster 4** | 9 | 0 |
| **Cluster 5** | 0 | 4 |
| **Cluster 6** | 5 | 0 |
| **Cluster 7** | 10 | 0 |
| **Cluster 8** | 1 | 3 |
| **Cluster 9** | 11 | 0 |
| **Cluster 10** | 3 | 1 |
| **Cluster 11** | 6 | 0 |
| **Cluster 12** | 3 | 0 |
| **Cluster 13** | 1 | 14 |
| **Cluster 14** | 0 | 5 |
| **Cluster 15** | 11 | 9 |
| **Cluster 16** | 5 | 4 |
| **Cluster 17** | 4 | 0 |
| **Cluster 18** | 1 | 2 |
| **Cluster 19** | 3 | 6 |
| **Cluster 20** | 2 | 2 |
| **Cluster 21** | 10 | 1 |
| **Cluster 22** | 2 | 4 |
| **Cluster 23** | 44 | 1 |
| **Cluster 24** | 14 | 0 |
| **Cluster 25** | 8 | 0 |
| **Cluster 26** | 0 | 3 |
| **Cluster 27** | 15 | 0 |
| **Cluster 28** | 3 | 3 |
| **Cluster 29** | 2 | 1 |
| **Cluster 30** | 0 | 8 |
| **Cluster 31** | 8 | 0 |
| **Unclustered ($N$ < 3)** | 670 | 53 |
